# Cell lines of the same anatomic site and histologic type show large variability in radiosensitivity and relative biological effectiveness to protons and carbon ions

**DOI:** 10.1101/2020.06.19.161497

**Authors:** David B. Flint, Scott J. Bright, Conor H. McFadden, Teruaki Konishi, Daisuke Ohsawa, Broderick Turner, Steven H. Lin, David R. Grosshans, Simona F. Shaitelman, Hua-Sheng Chiu, Pavel Sumazin, Gabriel O. Sawakuchi

**Affiliations:** The University of Texas MD Anderson Cancer Center UTHealth Graduate School of Biomedical Sciences, Houston, TX, USA; Department of Radiation Physics, The University of Texas MD Anderson Cancer Center, Houston, TX, USA; Single Cell Radiation Biology Group, Institute for Quantum Life Science, National Institutes for Quantum and Radiological Science and Technology, Inage-ku, Chiba, Japan; Department of Basic Medical Sciences for Radiation Damages, National Institute of Radiological Sciences, National Institutes for Quantum and Radiological Science and Technology, Inage-ku, Chiba, Japan; Department of Accelerator and Medical Physics, National Institute of Radiological Sciences, National Institutes for Quantum and Radiological Science and Technology, Inage-ku, Chiba, Japan; Department of Radiation Oncology, The University of Texas MD Anderson Cancer Center, Houston, TX, USA; Texas Children’s Cancer Center, Baylor College of Medicine

**Author notes:** Corresponding author: Gabriel O. Sawakuchi, PhD, Department of Radiation Physics, Unit 1420, The University of Texas MD Anderson Cancer Center, 6565 MD Anderson Blvd, Houston, TX 77030-4008.; email: g.

**Keywords:** relative biological effectiveness, radiosensitivity, particle therapy

## Abstract

**Purpose:** To show that radiation response across cancer cell lines of the same anatomic site and histologic type varies remarkably for protons and carbon (C) ions.

**Materials and Methods:** We measured and obtained from the literature clonogenic survival of human cancer cell lines of the lung (n=18), brain (n=10) and pancreas (n=10) exposed to photons, protons, and C-ions to assess their variability in response. We also treated cancer cell lines with DNA repair inhibitors prior to irradiation to assess how DNA repair capacity affects their variability in response. We quantified the variability in response by calculating the relative range (range/mean) and the coefficient of variation (COV) of the dose at 10% survival fraction (D_10%_) and relative biological effectiveness (RBE_10%_).

**Results:** The relative range of D_10%_ for lung cancer cell lines varied from 55-92% for photons, protons, and C-ions, with the relative range in RBE varying from 16-45% for protons and C-ions. For brain and pancreatic cancer cell lines, the relative range of D_10%_ varied from 95-112%, and 39-75%, respectively, with the relative range in RBE varying from 27-33% and 25-50%, respectively. However, the COVs in D_10%_ were approximately equal across radiation qualities, varying from 0.24±0.07–0.35±0.10, 0.35±0.09–0.69±0.62 and 0.13±0.03– 0.21±0.04 for lung, brain and pancreatic cancer cell lines, respectively. Greater relative ranges in D_10%_ were observed in the cell lines with inhibited DNA repair, varying from 108%-157% for photons, protons, and C-ions, with relative ranges in RBE varying from 29-67%. The COVs in the D_10%_ were also greater for the cell lines treated with inhibitors of DNA repair, varying from 0.34±0.09–0.41±0.06.

**Conclusions:** Cell lines of the same anatomic site and histologic type have a remarkable variability in response, not only to photons but also to protons and C-ions. We attributed this variability to differences in DNA repair capacity.

**Category:** Biological Physics and Response Prediction

## INTRODUCTION

Although personalized treatments are ideally customized to the biology and genotype of a tumor, decisions for most tumors regarding the radiotherapy (RT) type (photons, protons, or carbon ions) and the dose-fractionation regimen are made independent of a tumor’s particular biology or genotype, resulting in a “one-size-fits-all” approach based on histology, patient performance status and disease stage. This is further compounded in the case of heavy ion therapy, where the relative biological effectiveness (RBE) varies greatly across the treatment volume^1^ and is accounted for through the use of models such as the local effect model (LEM)^2-4^, or the microdosimetric kinetic model (MKM)^5,6^ which, in clinical practice, predict the RBE for different histologic types based on the response of a discrete number of cell lines in a small subset of histological types.^1,7-11^ This may explain, in part, the heterogeneous response of tumors to RT in any given cohort of patients.^12-15^

While an oft-cited advantage of heavy ions is their lesser dependence on tumor biology^16,17^, which in turn might mitigate this heterogeneous response, in this letter we show that even cancer cell lines of the same anatomic site and histologic type have remarkable differences in response to photons, protons, and even C-ions. We attribute these differences to variations in genomic features of cell lines including gene expression and mutations. This suggests that underlying genetic variations between tumors from different patients (and even within the same tumor) are important drivers of radiation response even for heavy ion therapy and should be accounted for as part of radiation treatment planning.

## MATERIALS AND METHODS

### Cell line panel

We used human cancer cell lines of three anatomical sites including lung (n=18), brain (n=10) and pancreas (n=10), obtained from literature sources including the Particle Irradiation Data Ensemble (PIDE) database^18^, Liu et al. 2015 ^19^ and Suzuki et al. 2000^20^ as well as collected by us (Table 1). The lung cancer cell lines were all non-small cell lung carcinoma (NSCLC), which were further classified in three histologic subtypes including lung adenocarcinoma (LUAD) (n=13), lung squamous cell carcinoma (LUSC) (n=4) and large cell lung carcinoma (LCLC) (n=1). However, no distinction between these subtypes is made in clinical practice to prescribe radiotherapy. NSCLC were then further classified according to their molecular subtypes, which were obtained from Yu et al. 2019.^21^ Brain cancer cell lines were all glioblastoma multiforme (GBM) (n=10) and were classified by molecular subtype according to PTEN-wildtype (n=4) or PTEN-mutant (n=6). All brain cancer cell lines were IDH-wildtype. PTEN and IDH mutation statuses were from the Cancer Cell Line Encyclopedia (CCLE).^22^ Notably, the GBM cell lines M059K and M059J were extracted from the same tumor but M059K has normal DNA-dependent protein kinase, catalytic subunit (DNA-PKcs) activity while M059J lacks DNA-PKcs activity. DNA-PKcs is essential for non-homologous end-joining repair (NHEJ) of double strand breaks. The pancreatic cancer cell lines were all pancreatic adenocarcinoma (PAAD), further classified in molecular subtypes including basal (n=5) and classical (n=5).^21^

**Table 1.**
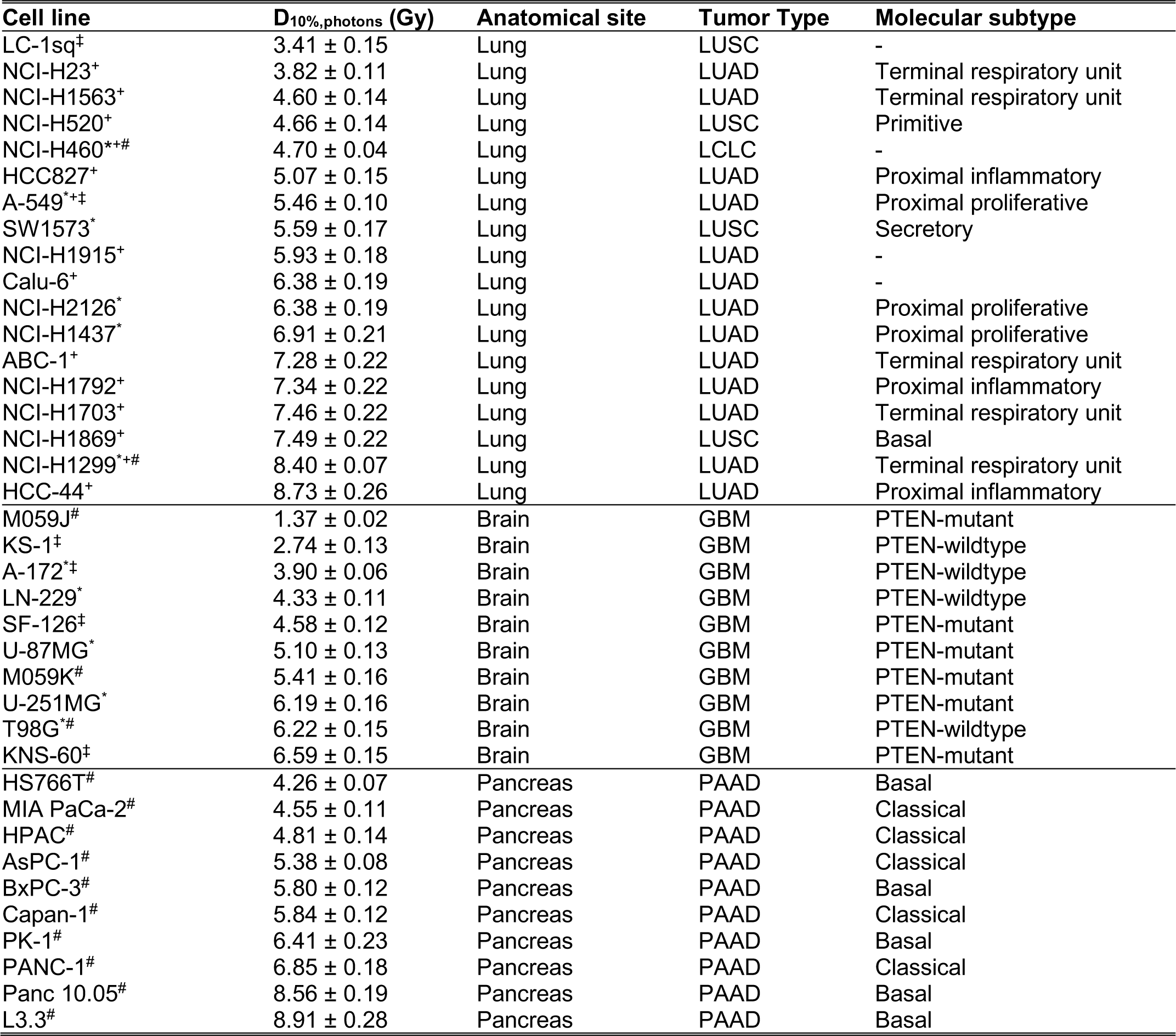
List of lung, brain and pancreatic cancer cell lines with their respective histology in order of radiosensitivity to photons. “^*^”, “^+^”, “^‡^” and “^#^” indicate D_10%,photon_ data from the PIDE database, Liu et al. 2015^19^, Suzuki et al. 2000^20^, or our measurements, respectively (multiple symbols indicates averaged from multiple sources). LUAD: Lung adenocarcinoma. LUSC: Lung squamous cell carcinoma. LCLC: Large cell lung carcinoma. GBM: Glioblastoma multiforme. PAAD: Pancreatic adenocarcinoma.

Additionally, H1299, H460, Panc 10.05 and PANC-1 cells were treated with DNA repair inhibitors prior to irradiation to attenuate various DNA repair pathways to differing degrees. These inhibitors targeted the proteins: ataxia telangiectasia and Rad3 related (ATR), DNA-PKcs, ataxia telangiectasia mutated (ATM), Poly (ADP-ribose) polymerase (PARP) and Rad51, which are proteins essential for DNA damage response (DDR); NHEJ repair, DDR, DDR and base excision repair (BER), and homologous recombination (HR) repair, respectively. Details of these drug treatments can be found below. These cell lines were selected because they represent two anatomic sites and for each anatomic site one cell line is radiosensitive and the other is radioresistant (Table 1).

### Treatment with DNA repair inhibitors

To inhibit ATR, Ceralasertib (AZD6738, Selleckchem) was dissolved in DMSO at 10 mM and was used at final concentrations of 0.1 or 2 µM. Alternatively to inhibit ATR, BAY 1895344, 2-[(3R)-3-methylmorpholin-4-yl]-4-(1-methyl-1H-pyrazol-5-yl)-8-(1H-pyrazol-5-yl)-1,7-naphthyridine (Selleckchem) was dissolved in DMSO at 10 mM and was used at a final concentration of 0.01 µM. To inhibit DNA-PKcs, 8-(4-Dibenzothienyl)-2-(4-morpholinyl)- 4H-1-benzopyran-4-one (NU7441, Selleckchem) was dissolved in DMSO at 5 mM and was used at final concentrations ranging from of 0.1 or 0.5 µM. To inhibit ATM, KU55933, 2-morpholin-4-yl-6-thianthren-1-yl-pyran-4-one (S1092, Selleckchem) was dissolved in DMSO at 10 mM and was used at final concentrations of 0.1 or 10 µM. To inhibit PARP1, Olaparib (AZD2281, S1060, Selleckchem) was dissolved in DMSO at 10 mM and was used at a final concentration of 1 µM. To inhibit Rad51, B02 (Cat# S8434, Selleckchem) was dissolved in DMSO at 10 mM and used at a final concentration of 5 μM. The final concentration of DMSO was 0.1% in all groups. See Table 2 for more details. Cells were seeded 24 hours prior to irradiation in 6-well plates or T12.5 flasks. Then, 8 hours prior to irradiation, media was removed and replaced with media containing inhibitor or DMSO vehicle to serve as control. After 24 hours of incubation with the inhibitor or vehicle (16 hours after irradiation), media was removed and replaced with fresh media. Cells were not washed to minimize disturbance to attached cells.

**Table 2.** List of lung and pancreatic cancer cell lines and their respective tumor types treated with varying concentrations of DNA repair inhibitors to modulate their radiosensitivity to photons. LUAD: Lung adenocarcinoma. LUSC: Lung squamous cell carcinoma. LCLC: Large cell lung carcinoma. PAAD: Pancreatic adenocarcinoma. DNA-PKcsi: cells treated with DNA-PKcs inhibitor Nu7441. ATMi: cell treated with ATM inhibitor KU55933. ATRi: cells treated with ATR inhibitor AZD6738. ATRi-BAY: cells treated with ATR inhibitor BAY 1895344. PARP1i: cells treated with PARP1 inhibitor Olaparib (AZD2281). RAD51i: cells treated with Rad51 inhibitor B02. DMSO: cell lines treated with DMSO vehicle as control. [] concentration of inhibitor.

| Cell line | D<sub>10%,photons</sub> (Gy) | Anatomical site | Tumor Type | Molecular subtype |
| --- | --- | --- | --- | --- |
| H1299+DNA-PKcsi [1μM] | 2.16 ± 0.25 | Lung | LUAD | Terminal respiratory unit |
| H460+DNA-Pkcsi [1 μM] | 2.34 ± 0.05 | Lung | LCLC | - |
| H460+ATMi [10 μM] | 2.38 ± 0.07 | Lung | LCLC | - |
| H460+ATRi [0.5 μM] | 2.66 ± 0.23 | Lung | LCLC | - |
| H460+DNA-PKcsi [0.5μM] | 2.70 ± 0.09 | Lung | LCLC | - |
| H1299+ATMi [10 μM] | 2.76 ± 0.10 | Lung | LUAD | Terminal respiratory unit |
| H1299+DNA-Pkcsi [0.5 μM] | 2.91 ± 0.10 | Lung | LUAD | Terminal respiratory unit |
| H460+PARP1i [1 μM] | 3.15 ± 0.07 | Lung | LCLC | - |
| H460+ATRi-BAY [0.01 μM] | 3.53 ± 0.07 | Lung | LCLC | - |
| H460+ATRi [0.1 μM] | 3.69 ± 0.06 | Lung | LCLC | - |
| H460+ATMi [0.1 μM] | 3.97 ± 0.09 | Lung | LCLC | - |
| H460+DNA-PKcsi [0.1 μM] | 3.98 ± 0.06 | Lung | LCLC | - |
| H460+RAD51i [5 μM] | 4.52 ± 0.04 | Lung | LCLC | - |
| H460+DMSO | 5.32 ± 0.03 | Lung | LCLC | - |
| H1299+ATRi – BAY [0.01 μM] | 6.12 ± 0.11 | Lung | LUAD | Terminal respiratory unit |
| H1299+PARP1i [1 μM] | 6.19 ± 0.10 | Lung | LUAD | Terminal respiratory unit |
| H1299+ATRi [0.5 μM] | 6.69 ± 0.35 | Lung | LUAD | Terminal respiratory unit |
| H1299+DNA-Pkcsi [0.1 μM] | 7.25 ± 0.13 | Lung | LUAD | Terminal respiratory unit |
| H1299+ATRi [0.1 μM] | 7.91 ± 0.33 | Lung | LUAD | Terminal respiratory unit |
| H1299+ATMi [0.1 μM] | 8.24 ± 0.23 | Lung | LUAD | Terminal respiratory unit |
| H1299+RAD51i [5 μM] | 8.71 ± 0.18 | Lung | LUAD | Terminal respiratory unit |
| H1299+DMSO | 8.91 ± 0.13 | Lung | LUAD | Terminal respiratory unit |
| PANC-1+ATMi [10 μM] | 2.83 ± 0.12 | Pancreas | PAAD | Classical |
| Panc 10.05+DNA-Pkcsi [0.5 μM] | 2.98 ± 0.15 | Pancreas | PAAD | Basal |
| Panc 10.05+ATMi [10 μM] | 3.01 ± 0.08 | Pancreas | PAAD | Basal |
| Panc 10.05+ATRi [2 μM] | 3.54 ± 0.18 | Pancreas | PAAD | Basal |
| PANC-1+DNA-Pkcsi [0.5 μM] | 3.59 ± 0.08 | Pancreas | PAAD | Classical |
| PANC-1+PARP1i [1 μM] | 4.29 ± 0.13 | Pancreas | PAAD | Classical |
| PANC-1+ATRi [2 μM] | 5.17 ± 0.15 | Pancreas | PAAD | Classical |
| Panc 10.05+PARP1i [1 μM] | 5.58 ± 0.10 | Pancreas | PAAD | Basal |
| Panc 10.05+ATRi [0.1 μM] | 5.75 ± 0.27 | Pancreas | PAAD | Basal |
| PANC-1+DNA-Pkcs [0.1 μM] | 5.79 ± 0.11 | Pancreas | PAAD | Classical |
| PANC-1+RAD51i [5 μM] | 5.87 ± 0.10 | Pancreas | PAAD | Classical |
| Panc 10.05+DNA-PKcs [0.1 μM] | 6.04 ± 0.15 | Pancreas | PAAD | Basal |
| PANC-1+ATMi [0.1 μM] | 6.10 ± 0.10 | Pancreas | PAAD | Classical |
| Panc 10.05+RAD51i [5 μM] | 6.24 ± 0.13 | Pancreas | PAAD | Basal |
| PANC-1+ATRi [0.1 μM] | 6.24 ± 0.11 | Pancreas | PAAD | Classical |
| PANC-1+DMSO | 6.26 ± 0.12 | Pancreas | PAAD | Classical |
| Panc 10.05+ATMi [0.1 μM] | 6.52 ± 0.19 | Pancreas | PAAD | Basal |
| Panc 10.05+DMSO | 7.42 ± 0.24 | Pancreas | PAAD | Basal |

### Radiation response data

Radiation response data of each cell line were either measured by us (H460, H1299, BxPC-3, PANC-1, AsPC-1 and Panc 10.05, MIA PaCa-2, L3.3, M059K and M059J cell lines) or obtained from the literature. The literature data included data extracted from the PIDE database^18^, from Liu et al. 2015^19^ and from Suzuki et al. 2000^20^. To quantify the radiosensitivity, the dose at which the surviving fraction falls to 10% (D_10%_) for the cell survival curve was calculated based on α and β values from the linear quadratic model, which were fitted to the clonogenic cell survival data or those provided in the literature sources. In cases where data existed from multiple sources, we took the average D_10%_ value. We quantified the ion relative biological effectiveness (RBE) relative to the photons by taking the ratio of D_10%_ values: RBE_D10%_ = D_10%,photon_/D_10%,ion_.

To quantify the variability of the response within each group, we calculated the range of the D_10%_ and RBE_D10%_ values. To compare between histologic types, ion linear energy transfer (LET) values, or treatment with DNA repair inhibitors, we calculated the relative range (range/mean) and the coefficient of variation (COV) of D_10%_ or RBE _D10%_.

### Cell culture

M059K, M059J, H460, H1299, BxPC-3, PANC-1, AsPC-1 and Panc 10.05, MIA PaCa-2 and L3.3 cell lines were obtained from the American Type Culture Collection (ATCC). Cell culture conditions are specified in the Supplementary Note 1. All cell lines were maintained in a humidified atmosphere at 37°C, 5% CO_2_ in air. All cell lines were routinely sub-cultured prior to reaching 100% confluence using 0.25% trypsin.

### Irradiations

Cell lines were grown in 6 well plates or T-12.5 flasks and exposed to clinical x-ray and proton beams at the Proton Therapy Center MD Anderson Cancer Center, and C-ions at the Heavy-Ion Medical Accelerator in Chiba (HIMAC) (Chiba, Japan). X-ray irradiations were done using a 6 MV beam from a clinical linac (Truebeam, Varian Medical Systems, Palo Alto, CA) at a water equivalent depth of 10 cm to simulate a clinical scenario. Proton irradiations were done using an unmodulated 100 MeV proton beam with slabs of water equivalent plastic in front of the beam to obtain a dose-weighted LET in water of 11.1 keV/μm, which corresponds to the end of the proton range. C-ion irradiations were done using unmodulated 290 MeV/nucleon C-ions with dose-weighted LET in water of 13.5 and 60.5 keV/μm, respectively, which correspond approximately to LET values found at the entrance and middle of SOBP for C-ions, respectively. These LET values were obtained using energy absorbers that are part of the beamline.^23^ Supplementary Note 2 provides detailed conditions of the irradiations.

### Clonogenic cell survival assay

Cells were trypsinized 18-24 hours prior to irradiation and seeded into 6-well plates or T-12.5 flasks at appropriate numbers for each dose. Cell lines were allowed to form colonies for 7-14 days and then were fixed and stained with pure ethanol containing 0.5% crystal violet. Plates were allowed to air dry overnight and then scanned using a high-resolution flatbed scanner (Epson Expression 10000 XL). Images were analyzed using an ImageJ plugin developed in-house that automatically counts colonies containing more than 50 cells.^24^ Details of the clonogenic cell survival assay are in Supplementary Note 3.

## Statistical analyses

Statistical analyses were performed in MATLAB 2017 (Mathworks, Natick, MA) and Graph Pad Prism 7 (Graph Pad, San Diego, CA). Error bars represent the standard error propagated from the survival curve parameters (α and β) fits, including their covariance, to the parameter estimates.

## RESULTS

We found large variability in the D_10%_ values of several lung, brain and pancreatic cancer cell lines to photons, protons, and even C-ions (Fig. 1A), varying 55±6–92±12%, 95±46–112±20% and 39±5–75±6% for lung, brain and pancreatic cancer cell lines, respectively, relative the mean of each group (Table 3). But when comparing between radiation qualities, while the absolute ranges in D_10%_ are much higher for photons than C-ions (Table 3), when D_10%_ for a particular radiation quality is normalized to the mean D_10%_ in each group, the relative D_10%_ values seemingly span similar ranges across all radiation qualities (Fig. 1B) (Table 3). Accordingly, when we quantify the variability in D_10%_ within each group by calculating the coefficient of variation (COV), no significant differences can be seen in the variability of D_10%_ between radiation qualities (Fig. 2A-C) (Table 3).

**Table 3:**
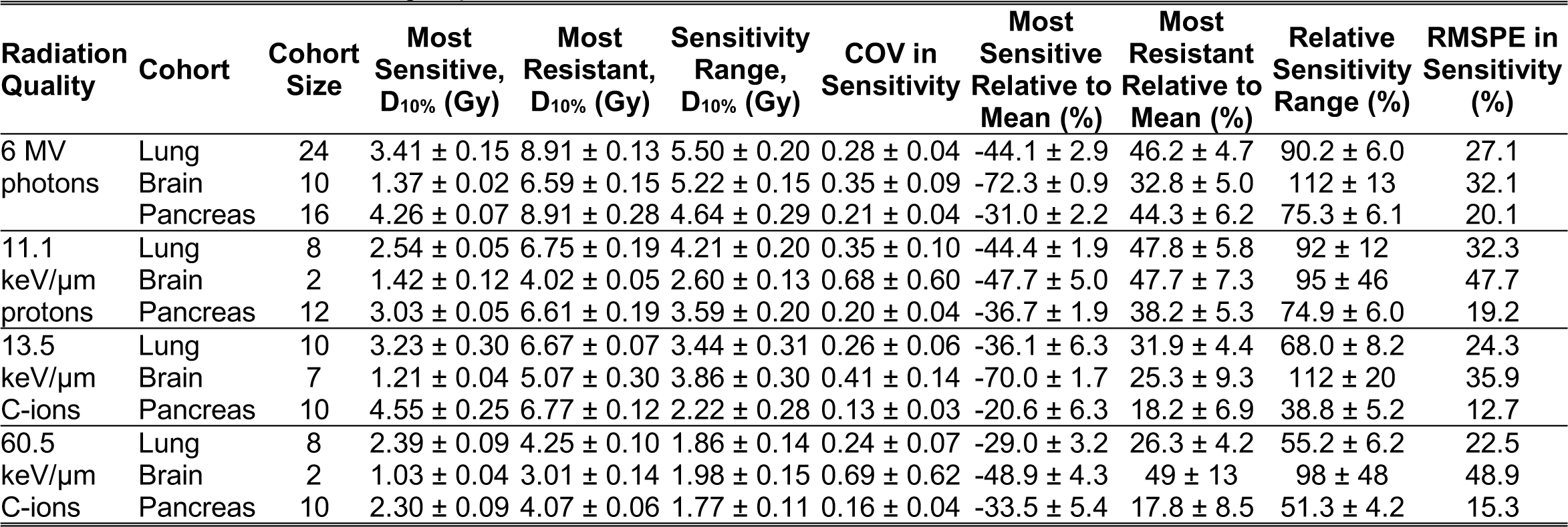
Variability in radiosensitivity of lung, brain and pancreatic cancer cells exposed to 6 MV photons, 11.1 keV/μm protons, and 13.5 and 60.5 keV/μm C-ions, grouped by cancer site. Relative sensitivity values are calculated relative to the mean D_10%_ value for each group.

| Radiation Quality | Cohort | Cohort Size | Most Sensitive, D<sub>10%</sub> (Gy) | Most Resistant, D<sub>10%</sub> (Gy) | Sensitivity Range, D<sub>10%</sub> (Gy) | COV in Sensitivity | Most Sensitive Relative to Mean (%) | Most Resistant Relative to Mean (%) | Relative Sensitivity Range (%) | RMSPE in Sensitivity (%) |
| --- | --- | --- | --- | --- | --- | --- | --- | --- | --- | --- |
| 6 MV photons | Lung | 24 | 3.41 ± 0.15 | 8.91 ± 0.13 | 5.50 ± 0.20 | 0.28 ± 0.04 | -44.1 ± 2.9 | 46.2 ± 4.7 | 90.2 ± 6.0 | 27.1 |
|  | Brain | 10 | 1.37 ± 0.02 | 6.59 ± 0.15 | 5.22 ± 0.15 | 0.35 ± 0.09 | -72.3 ± 0.9 | 32.8 ± 5.0 | 112 ± 13 | 32.1 |
|  | Pancreas | 16 | 4.26 ± 0.07 | 8.91 ± 0.28 | 4.64 ± 0.29 | 0.21 ± 0.04 | -31.0 ± 2.2 | 44.3 ± 6.2 | 75.3 ± 6.1 | 20.1 |
| 11.1 keV/μm protons | Lung | 8 | 2.54 ± 0.05 | 6.75 ± 0.19 | 4.21 ± 0.20 | 0.35 ± 0.10 | -44.4 ± 1.9 | 47.8 ± 5.8 | 92 ± 12 | 32.3 |
|  | Brain | 2 | 1.42 ± 0.12 | 4.02 ± 0.05 | 2.60 ± 0.13 | 0.68 ± 0.60 | -47.7 ± 5.0 | 47.7 ± 7.3 | 95 ± 46 | 47.7 |
|  | Pancreas | 12 | 3.03 ± 0.05 | 6.61 ± 0.19 | 3.59 ± 0.20 | 0.20 ± 0.04 | -36.7 ± 1.9 | 38.2 ± 5.3 | 74.9 ± 6.0 | 19.2 |
| 13.5 keV/μm C-ions | Lung | 10 | 3.23 ± 0.30 | 6.67 ± 0.07 | 3.44 ± 0.31 | 0.26 ± 0.06 | -36.1 ± 6.3 | 31.9 ± 4.4 | 68.0 ± 8.2 | 24.3 |
|  | Brain | 7 | 1.21 ± 0.04 | 5.07 ± 0.30 | 3.86 ± 0.30 | 0.41 ± 0.14 | -70.0 ± 1.7 | 25.3 ± 9.3 | 112 ± 20 | 35.9 |
|  | Pancreas | 10 | 4.55 ± 0.25 | 6.77 ± 0.12 | 2.22 ± 0.28 | 0.13 ± 0.03 | -20.6 ± 6.3 | 18.2 ± 6.9 | 38.8 ± 5.2 | 12.7 |
| 60.5 keV/μm C-ions | Lung | 8 | 2.39 ± 0.09 | 4.25 ± 0.10 | 1.86 ± 0.14 | 0.24 ± 0.07 | -29.0 ± 3.2 | 26.3 ± 4.2 | 55.2 ± 6.2 | 22.5 |
|  | Brain | 2 | 1.03 ± 0.04 | 3.01 ± 0.14 | 1.98 ± 0.15 | 0.69 ± 0.62 | -48.9 ± 4.3 | 49 ± 13 | 98 ± 48 | 48.9 |
|  | Pancreas | 10 | 2.30 ± 0.09 | 4.07 ± 0.06 | 1.77 ± 0.11 | 0.16 ± 0.04 | -33.5 ± 5.4 | 17.8 ± 8.5 | 51.3 ± 4.2 | 15.3 |

**Fig. 1.**
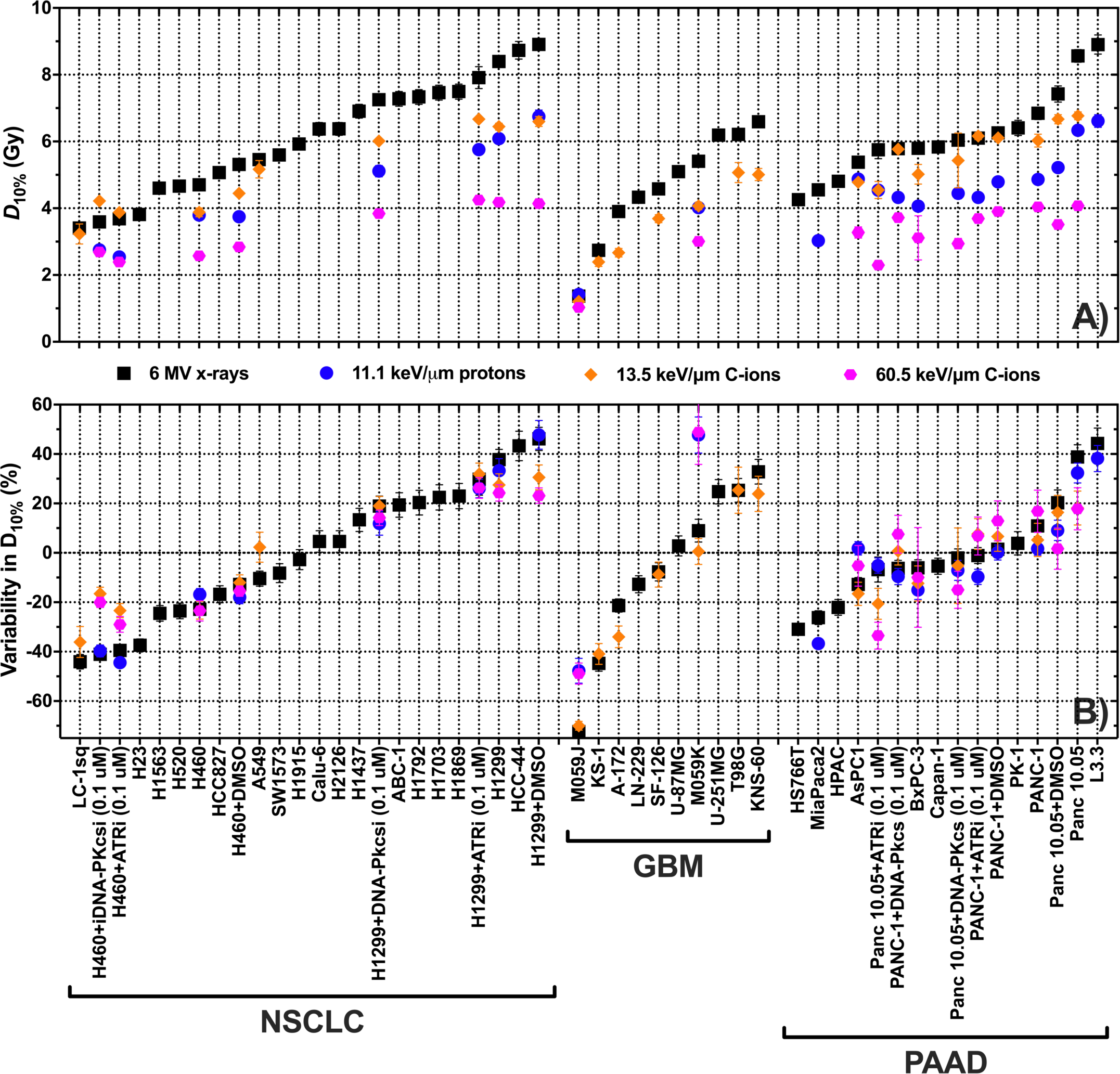
(A) Dose for 10% survival fraction to quantify survival from several NSCLC, GBM and PAAD cell lines. Each data point is the average between data we collected, data in PIDE database,^18^ data from Liu et al 2015,^19^ and data from Suzuki et al. 2000 ^20^, where available. (B) Percentage difference of the D_10%_ values from the mean D_10%_ for each radiation quality and histologic type to quantify the variation in response among cell lines of the same histologic type between different radiation qualities. NSCLC: Non-small cell lung carcinoma. GBM: Glioblastoma multiforme. PAAD: Pancreatic adenocarcinoma.

**Fig. 2:**
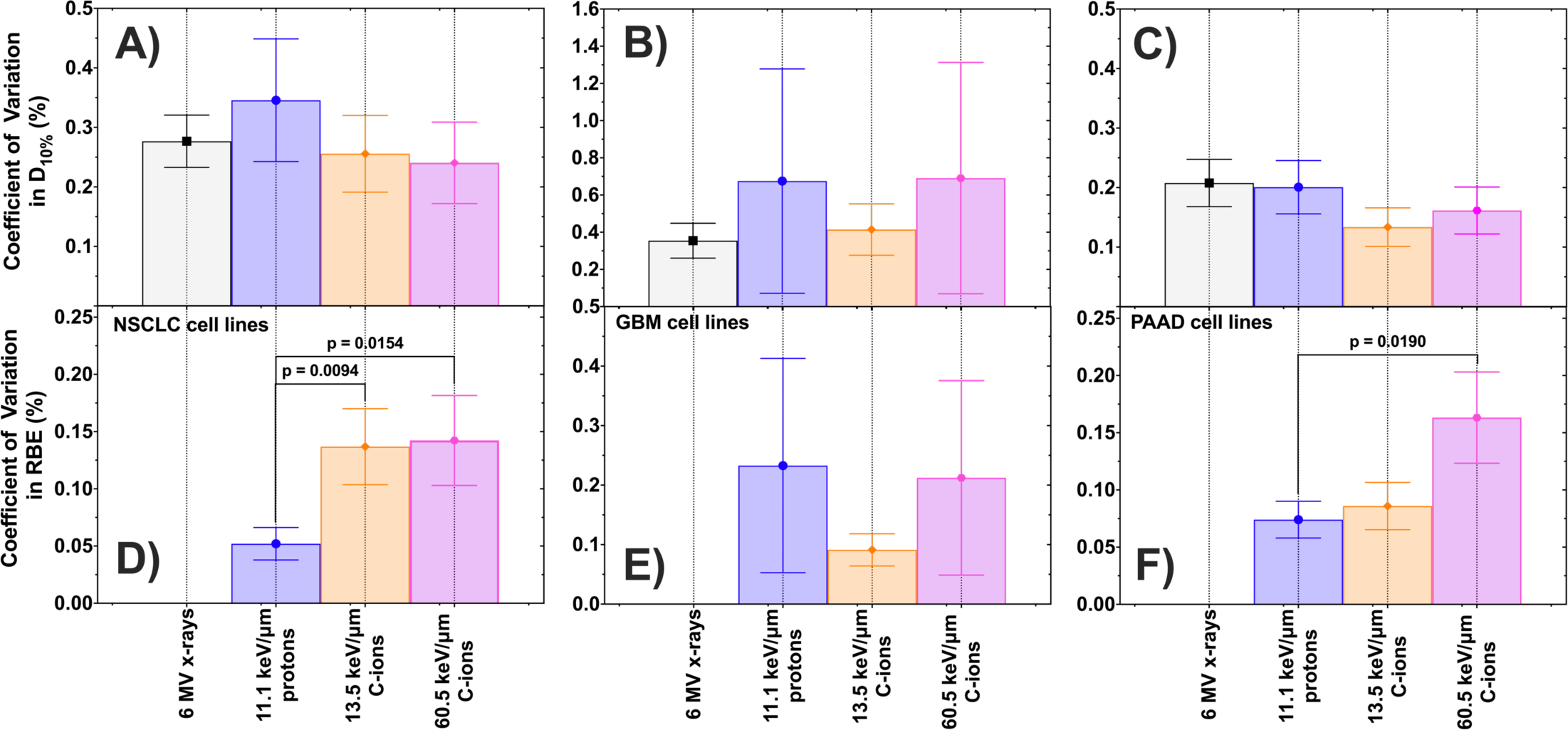
Coefficient of variation in D_10%_ (A-C) and RBE (D-F) of NSCLC (A,D), GBM (B, E) and PAAD (C,F) cancer cell lines.

Likewise, when the RBE values are calculated, we observe that there is large variability in the RBE _D10%_ values for protons and C-ions within each of the anatomic sites investigated (Fig. 3A), varying 16±3–45±4%, 26±7–33±10% and 25±9–50±9% for lung, brain and pancreatic cancer cell lines, respectively, relative the mean of each group (Table 4). But as with D_10%_, after normalizing the RBE _D10%_ for each radiation quality to the mean value in each anatomic site, we observe that the RBE _D10%_ values span similar relative ranges between radiation qualities (Fig. 3B) (Table 4). Moreover, when we calculate the COVs in RBE _D10%_, not only is the relative variability not significantly smaller for higher LET values, but in some cases, it is actually significantly higher for C-ions than for protons (P = 0.0154 and P = 0.0190 for lung and pancreatic cancer cells, respectively, exposed to 60.5 keV/μm C-ions) (Fig. 2D-F). Together, these data suggest that the relative variability in radiosensitivity and RBE are not smaller for higher LET radiation over the range of LET values investigated in this work.

**Table 4:** Variability in RBE of lung, brain and pancreatic cancer cells exposed to 11.1 keV/μm protons, and 13.5 and 60.5 keV/μm C-ions, relative to their 6 MV photon response, grouped by cancer site. Relative RBE values are calculated relative to the mean RBE_D10%_ value for each group.

| Radiation Quality | Cohort | Cohort Size | Lowest RBE | Highest RBE | RBE range | COV in RBE | Lowest RBE Relative to Mean (%) | Highest RBE Relative to Mean (%) | Relative RBE range (%) | RMSPE in RBE (%) |
| --- | --- | --- | --- | --- | --- | --- | --- | --- | --- | --- |
| 11.1 keV/μm protons | Lung | 8 | 1.24 ± 0.02 | 1.45 ± 0.04 | 0.21 ± 0.04 | 0.05 ± 0.01 | -9.1 ± 3.2 | 6.5 ± 4.4 | 16 ± 3 | 4.9 |
|  | Brain | 2 | 0.96 ± 0.08 | 1.34 ± 0.04 | 0.38 ± 0.09 | 0.23 ± 0.18 | -16.5 ± 9.8 | 16.5 ± 10.1 | 33 ± 10 | 16.5 |
|  | Pancreas | 12 | 1.10 ± 0.02 | 1.50 ± 0.05 | 0.40 ± 0.05 | 0.07 ± 0.02 | -18.4 ± 3.3 | 11.1 ± 5.4 | 29 ± 4 | 7.1 |
| 13.5 keV/μm C-ions | Lung | 10 | 0.85 ± 0.02 | 1.35 ± 0.04 | 0.50 ± 0.04 | 0.14 ± 0.03 | -25.1 ± 3.7 | 18.7 ± 6.2 | 43.8 ± 4.0 | 13.0 |
|  | Brain | 7 | 1.13 ± 0.04 | 1.46 ± 0.08 | 0.33 ± 0.09 | 0.09 ± 0.03 | -8.9 ± 5.9 | 17.8 ± 8.5 | 26.3 ± 7.0 | 8.8 |
|  | Pancreas | 10 | 0.99 ± 0.02 | 1.27 ± 0.09 | 0.27 ± 0.09 | 0.09 ± 0.02 | -11.5 ± 6.2 | 13.1 ± 11.1 | 24.6 ± 8.5 | 8.2 |
| 60.5 keV/μm C-ions | Lung | 8 | 1.34 ± 0.03 | 2.15 ± 0.04 | 0.81 ± 0.05 | 0.14 ± 0.04 | -26.2 ± 3.0 | 18.8 ± 4.5 | 45.0 ± 3.5 | 13.3 |
|  | Brain | 2 | 1.33 ± 0.06 | 1.80 ± 0.10 | 0.47 ± 0.12 | 0.21 ± 0.16 | -15.0 ± 7.2 | 15.0 ± 10.7 | 30.0 ± 8.6 | 15.0 |
|  | Pancreas | 10 | 1.56 ± 0.03 | 2.50 ± 0.16 | 0.95 ± 0.16 | 0.16 ± 0.04 | -17.2 ± 7.2 | 33.3 ± 14.0 | 50.4 ± 8.8 | 15.5 |

**Fig. 3.**
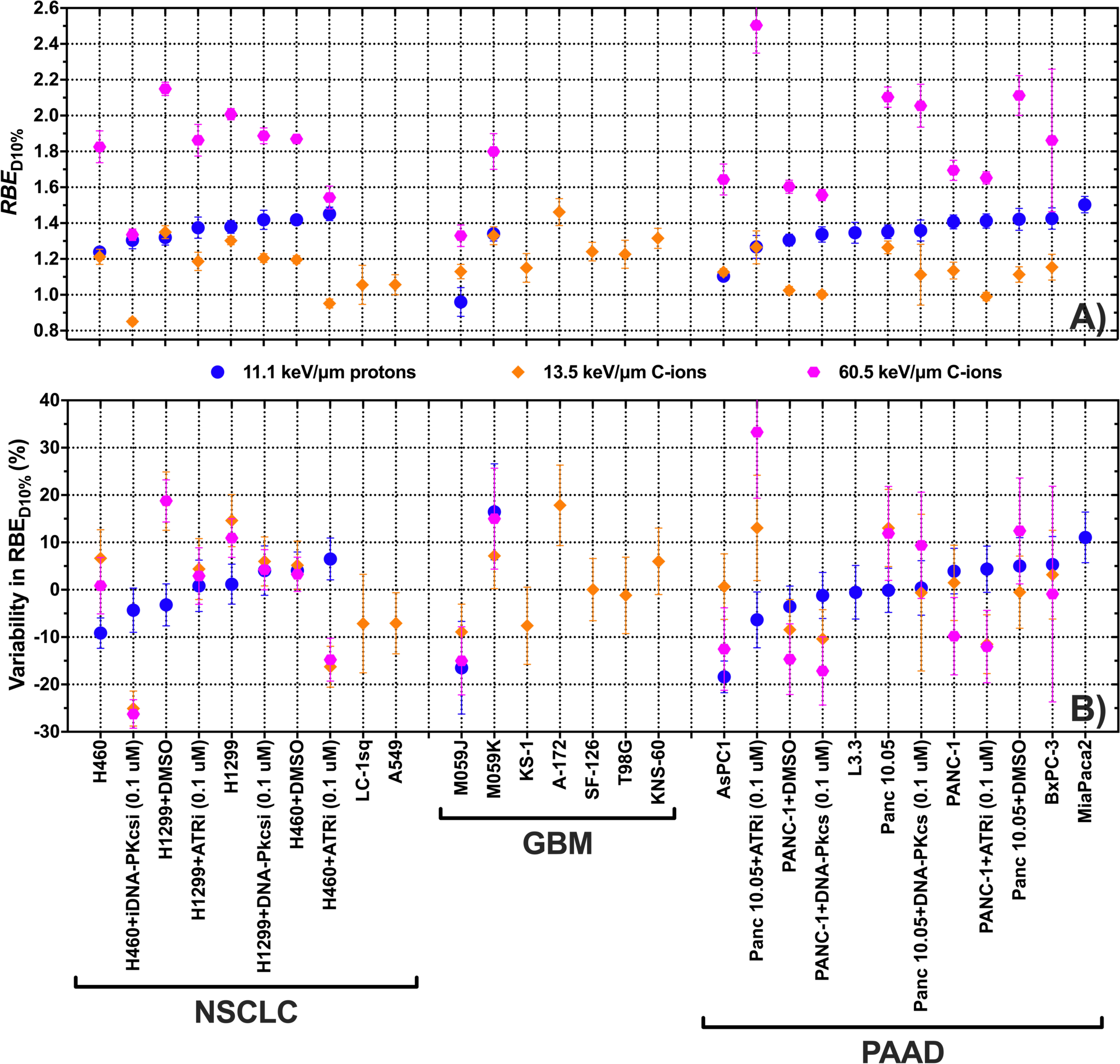
(A) RBE at the dose for 10% surviving fraction for NSCLC, GBM and PAAD cancer cell lines. Each data point is the average between data we collected, data in PIDE database ^18^, data from Liu et al 2015 ^19^, and data from Suzuki et al. 2000 ^20^, where available. (B) Percent difference in the RBE values relative to the mean RBE_D10%_ for each radiation quality and histologic type to quantify the variation in response among cell lines of the same histologic type between different radiation qualities.

Among the GBM cell lines examined, M059K and M059J are a naturally occurring isogenic pair extracted from the same tumor in which M059K has normal activity of DNA-PKcs while M059J lacks DNA-PKcs activity, rendering M059K 3.94 ± 0.13 times more radioresistant than M059J. To simulate this phenomenon across other histologic types, we treated cells with DNA repair inhibitors to reduce their capacity for DNA repair to varying degrees across different pathways, which also modulated their radiosensitivity (Fig. 4A). This resulted in a greater range of D_10%_ values in the cells with inhibited DNA repair (or a naturally occurring deficiency) than the uninhibited cells (Fig. 4A) (Table 5), a greater relative range in their D_10%_ values (Fig. 4B), and greater COVs for each radiation quality (Fig. 5A) (Table 5), with statistically significant differences in the COVs occurring for 6 MV photons (p = 0.0247), but with less significant differences for proton and C-ions (p = 0.08 – 0.23) (Fig. 5A).

**Table 5:** Variability in radiosensitivity of lung, brain and pancreatic cancer cells exposed to 6 MV photons, 11.1 keV/μm protons, and 13.5 and 60.5 keV/μm C-ions. Cells are grouped independent of histology but by whether the cells have been treated with DNA-repair inhibitors (or have a natural DNA-repair deficiency) versus cells whose DNA repair was not inhibited. Relative sensitivity values are calculated relative to the mean D_10%_ value for each group.

| Radiation Quality | Cohort | Cohort Size | Most Sensitive, D<sub>10%</sub> (Gy) | Most Resistant, D<sub>10%</sub> (Gy) | Sensitivity Range, D<sub>10%</sub> (Gy) | COV in Sensitivity | Most Sensitive Relative to Mean (%) | Most Resistant Relative to Mean (%) | Relative Sensitivity Range (%) | RMSPE in Sensitivity (%) |
| --- | --- | --- | --- | --- | --- | --- | --- | --- | --- | --- |
| 6 MV photons | Uninhibited DNA-repair | 9 | 4.70 ± 0.04 | 8.91 ± 0.13 | 4.21 ± 0.13 | 0.23 ± 0.06 | -31.5 ± 1.6 | 29.7 ± 3.5 | 61.2 ± 5.0 | 21.3 |
| | Inhibited DNA-repair | 37 | 1.37 ± 0.02 | 8.71 ± 0.18 | 7.34 ± 0.18 | 0.41 ± 0.06 | -70.6 ± 1.1 | 86.5 ± 7.2 | 157 ± 11 | 40.4 |
| 11.1 keV/μm protons | Uninhibited DNA-repair | 4 | 3.75 ± 0.07 | 6.75 ± 0.19 | 3.00 ± 0.20 | 0.22 ± 0.06 | -26.1 ± 2.1 | 33.1 ± 4.9 | 59.2 ± 5.9 | 20.9 |
| | Inhibited DNA-repair | 33 | 1.42 ± 0.12 | 6.62 ± 0.17 | 5.20 ± 0.21 | 0.36 ± 0.05 | -61.5 ± 3.4 | 79.3 ± 7.1 | 141 ± 10 | 35.0 |
| 13.5 keV/μm C-ions | Uninhibited DNA-repair | 9 | 3.88 ± 0.13 | 6.77 ± 0.12 | 2.89 ± 0.17 | 0.21 ± 0.06 | -31.6 ± 2.8 | 19.4 ± 3.5 | 51.0 ± 4.7 | 19.7 |
| | Inhibited DNA-repair | 9 | 1.21 ± 0.04 | 6.67 ± 0.07 | 5.45 ± 0.09 | 0.34 ± 0.10 | -75.1 ± 1.8 | 36.7 ± 8.8 | 112 ± 13 | 32.2 |
| 60.5 keV/μm C-ions | Uninhibited DNA-repair | 9 | 2.58 ± 0.12 | 4.18 ± 0.05 | 1.61 ± 0.13 | 0.17 ± 0.05 | -28.2 ± 4.0 | 16.6 ± 3.5 | 44.8 ± 4.6 | 16.4 |
| | Inhibited DNA-repair | 9 | 1.03 ± 0.04 | 4.25 ± 0.10 | 3.22 ± 0.11 | 0.34 ± 0.09 | -65.4 ± 1.7 | 42.4 ± 5.3 | 108 ± 13 | 31.8 |

**Fig. 4:**
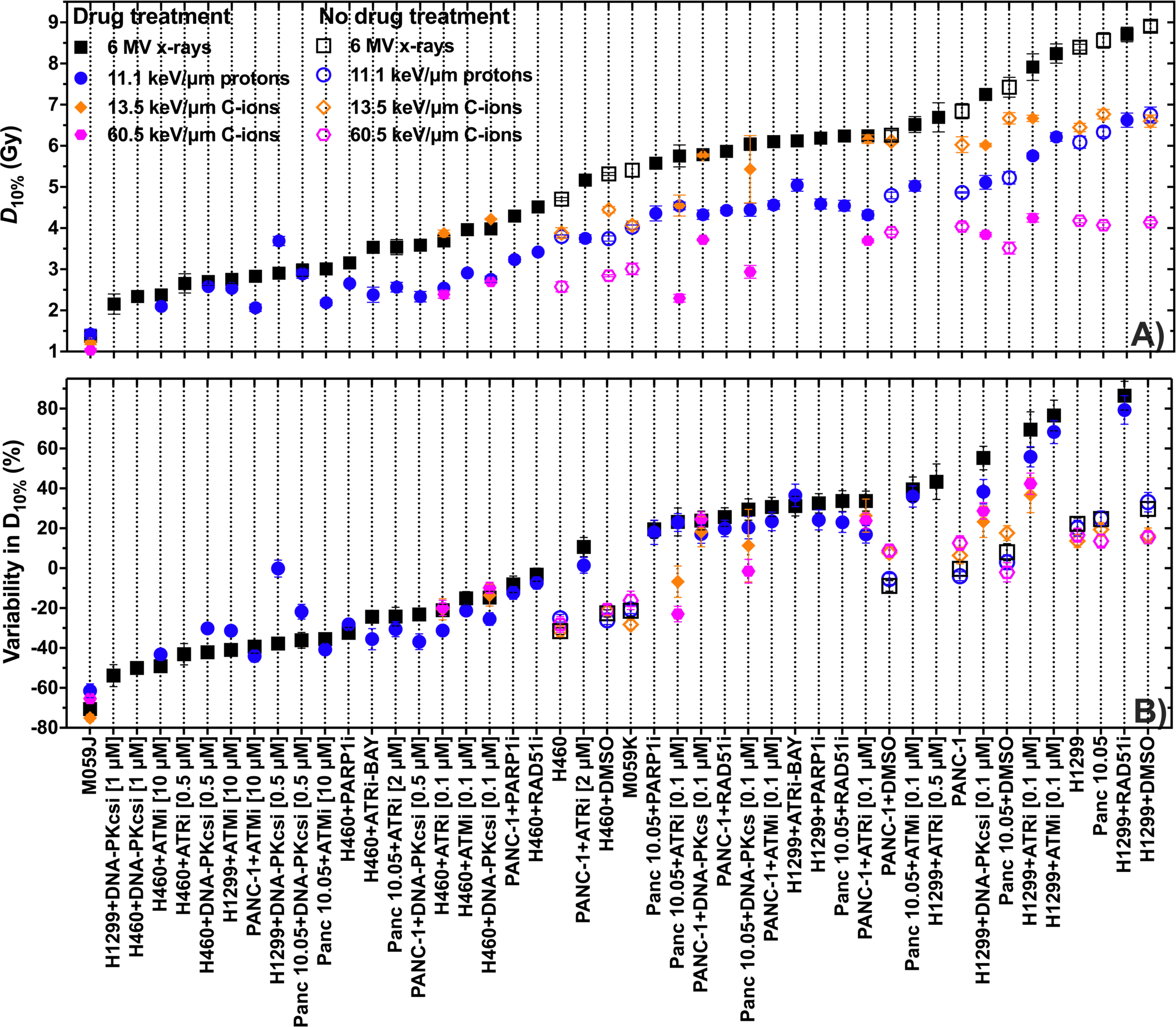
(A) Dose at 10% survival fraction for cell lines with DNA repair pathway defects or that were treated with DNA repair inhibitors (solid), or that have no defects and were not treated with inhibitors (open), exposed to 6 MV photons, 11.1 keV/µm protons, and 13.5 and 60.5 keV/μm C-ions. (B) Percentage difference relative to the mean values D10% value for each radiation quality and group to quantify the variation in response within groups between different radiation qualities.

**Fig. 5:**
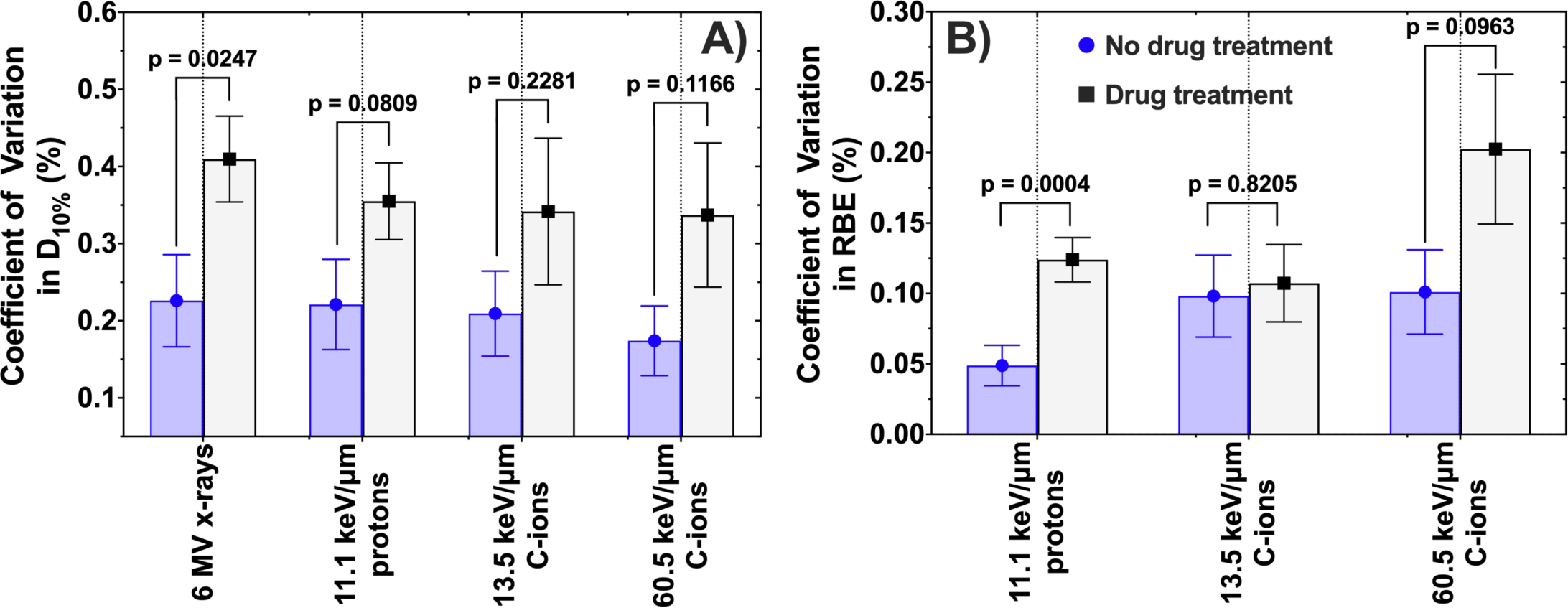
Coefficient of variation in (A) D_10%_ and (B) RBE of lung, brain and pancreatic cancer cells with DNA repair pathway defects or that were treated with DNA repair inhibitors (black), or that have no defects and were not treated with inhibitors (blue), exposed to 6 MV photons, 11.1 keV/µm protons, 13.5 keV/μm C-ions and 60.5 keV/μm C-ions, with the RBE values calculated relative to the 6 MV photon response.

The RBE _D10%_ values of the cells with inhibited DNA repair also span a wider range than the uninhibited cells (Fig. 6A), and there is also a greater range in their relative variation (Fig. 6B) (Table 6). In calculating the COVs in their RBE _D10%_ values, we confirmed that there is significantly more variation in proton RBE values in the cells with inhibited DNA repair than in the uninhibited cells (P=0.0004), but that this difference is less significant for higher LET radiation (P = 0.0963 for 60.5 keV/μm) (Fig. 5B) (Table 6). Together, these data suggest that modulating a cell’s capacity for DNA repair can significantly increase variability in their sensitivity and RBE among cell lines, implying that variations in cellular DNA repair capacity may still strongly influence cellular radiation response even for high LET radiation.

**Table 6:** Variability in RBE of lung, brain and pancreatic cancer cells exposed to 11.1 keV/μm protons, and 13.5 and 60.5 keV/μm C-ions, relative to their 6 MV photon response. Cells are grouped independent of histology but by whether the cells have been treated with DNA-repair inhibitors (or have a natural DNA-repair deficiency) versus cells whose DNA repair was not inhibited. Relative RBE values are calculated relative to the mean RBE_D10%_ value for each group.

| Radiation Quality | Cohort | Cohort Size | Lowest $\text{RBE}_{\text{D10\%}}$ | Highest $\text{RBE}_{\text{D10\%}}$ | $\text{RBE}_{\text{D10\%}}$ Range | COV in $\text{RBE}_{\text{D10\%}}$ | Lowest $\text{RBE}_{\text{D10\%}}$ Relative to Mean (%) | Highest $\text{RBE}_{\text{D10\%}}$ Relative to Mean (%) | Relative $\text{RBE}_{\text{D10\%}}$ Range (%) | RMSPE in $\text{RBE}_{\text{D10\%}}$ (%) |
| --- | --- | --- | --- | --- | --- | --- | --- | --- | --- | --- |
| 11.1 keV/$\mu\text{m}$ protons | Uninhibited DNA-repair | 7 | $1.24 \pm 0.02$ | $1.42 \pm 0.06$ | $0.18 \pm 0.06$ | $0.05 \pm 0.01$ | $-8.0 \pm 3.1$ | $5.6 \pm 5.6$ | $13.6 \pm 6.4$ | 4.5 |
| | Inhibited DNA-repair | 33 | $0.79 \pm 0.04$ | $1.54 \pm 0.09$ | $0.75 \pm 0.10$ | $0.12 \pm 0.02$ | $-39.1 \pm 3.8$ | $18.8 \pm 8.7$ | $57.9 \pm 9.6$ | 12.2 |
| 13.5 keV/$\mu\text{m}$ C-ions | Uninhibited DNA-repair | 7 | $1.02 \pm 0.02$ | $1.35 \pm 0.04$ | $0.33 \pm 0.04$ | $0.10 \pm 0.03$ | $-15.9 \pm 3.2$ | $10.8 \pm 4.6$ | $26.7 \pm 5.6$ | 9.1 |
| | Inhibited DNA-repair | 9 | $0.94 \pm 0.02$ | $1.27 \pm 0.09$ | $0.32 \pm 0.09$ | $0.11 \pm 0.03$ | $-13.4 \pm 6.1$ | $16 \pm 12$ | $29 \pm 13$ | 10.1 |
| 60.5 keV/$\mu\text{m}$ C-ions | Uninhibited DNA-repair | 7 | $1.60 \pm 0.04$ | $2.15 \pm 0.04$ | $0.55 \pm 0.05$ | $0.10 \pm 0.03$ | $-16.0 \pm 3.8$ | $12.6 \pm 4.9$ | $28.6 \pm 6.2$ | 9.4 |
| | Inhibited DNA-repair | 9 | $1.33 \pm 0.06$ | $2.50 \pm 0.16$ | $1.18 \pm 0.16$ | $0.20 \pm 0.05$ | $-24.8 \pm 4.8$ | $42 \pm 11$ | $67 \pm 12$ | 19.1 |

**Fig. 6:**
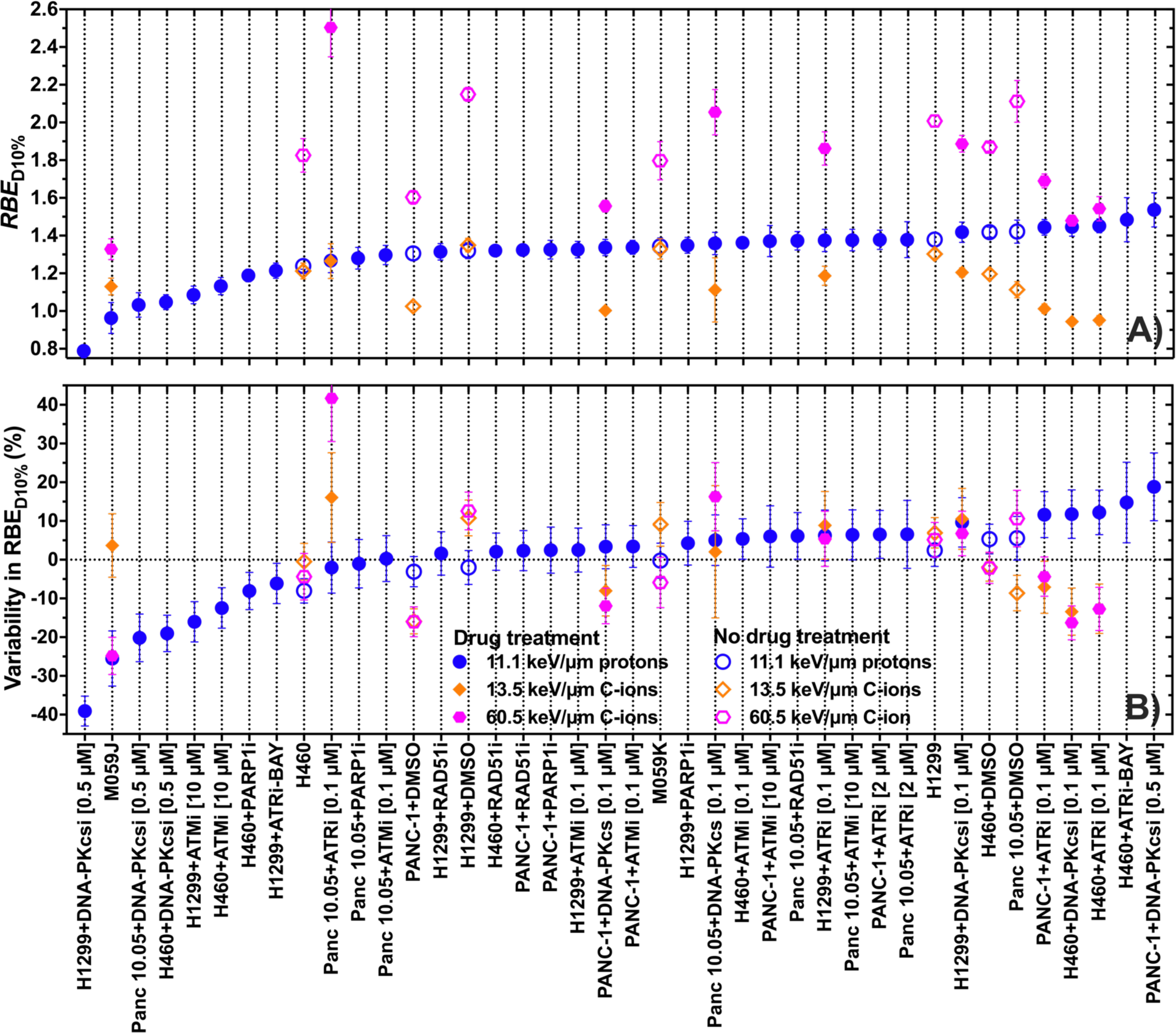
(A) RBE at the dose for 10% surviving fraction for lung, brain and pancreatic cancer cell lines with DNA repair pathway defects or that were treated with DNA repair inhibitors (solid), or that have no defects and were not treated with inhibitors (open), exposed to 11.1 keV/µm protons, and 13.5 and 60.5 keV/μm C-ions relative to their 6 MV photon response. (B) Percentage difference relative to the mean RBE_D10%_ value for radiation quality and group to quantify the variation in response within groups between different radiation qualities.

## DISCUSSION

Current practice in RT is to prescribe doses and fractionation schedules based on anatomic site, histology, patient’s performance status and disease staging, disregarding the tumor’s particular biology, genotype and the radiosensitization effect of any additional treatments. In particular, RT prescriptions for NSCLC, GBM and PAAD patients do not consider tumor subtype classification or molecular subtypes. But we showed that even cancer cell lines representing the same anatomic site and histologic type have remarkable differences in radiation response, not only when exposed to photons, but even in cells exposed to C-ions. This suggests that, across all RT modalities, for a given cohort of patients whose tumors are treated with the same fractionation schedule, current practice likely results in suboptimal treatments whose biological effects are potentially largely under- or overestimated. This may in turn contribute to poor outcomes, from either suboptimal tumor control or subjecting normal tissues to unnecessarily high doses.

We recently set out to test whether genomic variability can account for some of the observed variability in radioresistance between cell lines. Our results suggested the variability in gene expression, copy number variations, and mutation profiles can account for some of the variability in radiation response. Specifically, we found that the expression profiles of multiple genes that are known to play a role in DDR and DNA repair, are predictive of radiosensitivity and radioresistance. Here, we showed that radiosensitivity and radioresistance strongly depend on protein expression by inhibiting ATR, DNA-PKcs, ATM, PARP and Rad51, and by comparing isogenic cell lines with variable DNA-PKcs activity. Thus, our data suggested that genomic features of cell lines (and possibly tumors) are important factors influencing radiosensitivity and radioresistance of photons, protons and C-ions.

Furthermore, our data also indicated these genomic variations may drive radiation response not only to photons but also to higher LET radiation including protons and C-ions (Fig. 1, Fig. 4). Because the RBE _D10%_ values of the cell lines of the same anatomic site and histologic type vary greatly for a fixed LET of protons or C-ions (Fig. 3, Fig. 6), we argue that failing to incorporate these variabilities into the RBE estimate during treatment planning could further lead to unacceptable errors in RBE-weighted doses. Thus, we argue it is important to consider tumor genomic information into our treatment planning workflow and to predict RBE to individual patients and tumors rather than to a whole histologic type.

Furthermore, while RBE is an important parameter in heavy ion therapy, in the context of tumor-to-tumor variability, we argue the RBE itself cannot best be used without knowledge of radiosensitivity. This is because, as our data shows, cell lines of the same anatomic site and histologic type may have comparable RBE values while having a wide range of radiosensitivities. As illustrative examples, when the RBE values and relative D_10%,C-ions_ of cells exposed to 60.5 keV/μm C-ions are compared, the lung cancer cell lines H1299 and H460 have RBE values of 2.01 ± 0.03 and 1.83 ± 0.09, respectively, while D_10%,C-ions_ of H460 is 60% lower than that of H1299; the GBM cell lines M059K and M059J have RBE values of 1.80 ± 0.10 and 1.33 ± 0.06, respectively, while D_10%,C-ions_ of M059J is 190% lower than that of M059K; and the pancreatic cancer cell lines PANC-1 and AsPC-1 have RBE values of 1.69 ± 0.06 and 1.64 ± 0.09, respectively, while D_10%,C-ions_ of ASPC-1 is 20% lower than that of PANC-1 (Table 3 and Table 4). Thus, we argue that since the relative differences in D_10%_ can be much larger than the relative differences in RBE, RBE should be used in combination with radiosensitivity to allow proper dose prescriptions for individual tumors.

We believe that towards personalized RT, the consideration of genomic and tumor-microenvironment features has the potential to dramatically improve RT outcomes. These features include RNA, DNA, protein, and metabolome features, as well as features defining the hypoxic and immune infiltration status of the tumor. Ideally, these should be used in combination to inform clinicians on the radiosensitivity of individual tumors so that optimal fractionation schedules and choice of radiation type can be chosen rationally.

Moreover, as protons and C-ions are expensive and not available for all patients, it is important to select patients with radioresistant tumors who will most benefit from these radiation types. Personalized RT will depend on developing methods to predict tumor response to radiation prospectively and progress has been made in identifying genomic markers that may predict clinical response to photons.^25-31^ However, genomic models to predict tumor responses to protons and C-ions are not yet available. For optimal and rational use of the right technology – photons, protons or C-ions – for a given tumor, work should be focused on developing models able to predict tumor response not only to photons but also to protons and C-ions. In particular, the data presented in this work may be further analyzed to determine if molecular subtypes of each particular anatomic site better associate with radiation response. The data could also be used to further develop predictive models, including models based on DNA repair and DDR gene expression profiles, copy number variations or mutations.

A limitation of our current work is the reliance on literature data from different sources as well as the inclusion of cells treated with DNA repair inhibitors in the datasets used to quantify the variability for different radiation qualities. While this was necessary in order to have sufficient samples in a given group to quantify the variability within it to a reasonable level of precision, it may be viewed as artificially increasing the variability within the groups and thus confounding our conclusions. However, there are a number of reasons we believe that is not the case. First, we generally included more literature data for photons than ions due to the greater availability of photon data. Thus, if including literature data from different sources increased the variability, we would have expected to increase the variability more in the photon groups than in the ion groups. However, we did not see greater variability in the photon data than the C-ion data, so our conclusion that the C-ion data is no less variable than the photon data is not confounded. Second, when comparing between radiation qualities, we only included data of cells treated with DNA repair inhibitors for treatments that were performed across all radiation qualities and that induced radiosensitivities within the range of those established with naturally occurring cell lines. Thus any small additional variability this may have added is present across all the conditions we compared and should not confound our conclusions when comparing between them.

Furthermore, even if the inclusion of the cells treated with DNA repair inhibitors added different amounts of variability for different LET values, our secondary conclusions (Fig. 5A) show that, if anything, this should have added more variability to the 6 MV dataset than the ion datasets since differences between the inhibited versus uninhibited cohorts were most significant for this radiation quality. But this would have led us towards concluding that there is less variability in the ion dataset than the photon dataset while our data support the opposite conclusion: that the ion data is no less variable than photon data.

## CONCLUSIONS

Cell lines of the same anatomic site and histologic type vary remarkably in their response to photons, protons, and C-ions. We attributed this variability in radiation response to genomic differences across cell lines, perhaps in part due to differences in DNA repair capacity which are also expected to occur across tumors of the same anatomic site and histologic type. We also noted that the variability in radiation response has similar relative magnitudes across radiation types over the LET range investigated (up to 60.5 keV/μm) and is not smaller for higher LET values. We therefore conclude that it is important to develop predictive models of radiation response that incorporate the genomic variability observed among tumors, especially to allow proper selection of patients for particle therapy.

## ACKNOWLEDGMENTS

This work was supported in part by the University Cancer Foundation via the Sister Institution Network Fund (G.O.S.) and Institutional Research Grant program (G.O.S.) at the University of Texas MD Anderson Cancer Center; by the Cancer Prevention and Research Institute of Texas grant RP170040 (G.O.S.); and by the Cancer Center Support (Core) Grant 2P30CA016672-43 to MD Anderson. The authors also thank Dr. Narayan Sahoo at the MD Anderson Proton Therapy Center for proton beam-time scheduling at the MD Anderson Proton Center.

