## Supplemental Material for "Cell lines of the same anatomic site and histologic type show large variability in radiosensitivity and relative biological effectiveness to protons and carbon ions"

**Supplementary Note 1: Cell lines and details on culturing conditions**

The technical details of the cell culture media and additives are given in Supplemental Table 1.
The cells were cultured in T-25 and T-75 flasks. 24 h prior to irradiation, they were seeded into either 6-well plates or T-12.5 flasks to perform the clonogenic assays. To transfer the cells from the culture flasks to the colongenic vessels, the cells were washed with phosphate buffered saline (PBS), and treated with trypsin. For the exposures at the HIMAC's horizontal beamline, 2 hours prior to irradiation the 6-well plates filled with ~15 ml of media and sealed with aluminum sealers to ensure complete media of each well. The sealers were perforated immediately after irradiation to restore airflow. Details of these reagents and equipment are given in Supplemental Table 2.

Supplemental Table 1: Cell culture media and additives.

| Cell Line | Culture Medium; Details – Manufacturer (Cat. #) | Media Additives – Manufacturer (Cat. #) |
|---|---|---|
| M059K<br>M059J | 1:1 DMEM/F-12 Ham; with L-glutamine, NaHCO3, without HEPES – Sigma (D8062) | 10% FBS – Sigma (F0926)<br>1% PS – Hyclone (SV30010)<br>15 mM HEPES – Sigma (H0887) |
| H1299<br>H460<br>BxPC-3 | RPMI 1640; with L-glutamine, NaHCO3 – Sigma (R8758) | 10% FBS – Sigma (F0926)<br>1% PS – Hyclone (SV30010) |
| AsPC-1<br>PANC-1<br>Panc 10.05 | DMEM: high glucose; with 4500 mg/L glucose, L-glutamine, sodium pyruvate, NaHCO3 – Sigma (D6429) | 10% FBS – Sigma (F0926)<br>1% PS – Hyclone (SV30010 |

Supplemental Table 2: Reagents and supplies used to culture and process the cells.

| Reagent/Supply | Manufacturer (Cat. #) |
|---|---|
| T-25 flask | Thermo Scientific (156367) |
| T-75 flask | Thermo Scientific (156499) |
| T-12.5 flask | Cell Treat (229321) |
| 6-well plates | Corning Costar (3506) |
| PBS | Hyclone (SH30256.01) |
| Trypsin | Corning (25-053-CL) |
| Aluminum Sealer | Bio Rad (MSF1001) |
| Crystal Violet | Sigma (C0775-25G) |

### Supplementary Note 2: Irradiation details and dosimetry

#### *X-ray irradiations*

X-ray irradiations at MD Anderson were done using a clinical 6 MV x-ray flat beam (Truebeam, Varian Medical Systems, Palo Alto, CA) at water equivalent depth of 10 cm, which was obtained using slabs of water equivalent material in front of the beam. To avoid air gaps and provide lateral scattering and backscattering conditions we used a 6-well holder that can hold four 6-well plates. At least 9 cm of water equivalent slabs were placed downstream the plates to provide full backscattering conditions. Dummy plates with wells pre-filled with water by the same volume of medium were placed in the irradiation beam if less than 4 plates were exposed at the same time. Plates were exposed at 180º gantry angle at maximum dose rate (600 MU/min) and beam traversed the couch, water equivalent slabs and bottom of the plate before reaching the cells. Field size were 40 cm × 40 cm. Dosimetry in the same geometrical conditions were performed using small optically stimulated luminescence dosimeters (OSLD) placed inside of the wells of the 6-well plates.

#### *Proton irradiations*

Proton irradiations at MD Anderson were performed using an unmodulated 100 MeV proton beam (range 4.3 cm) at water equivalent depth of 4.4 cm, which was obtained using slabs of water equivalent material in front of the beam. To obtain the unmodulated beam, the range modulator wheel was parked in its thinnest step. Irradiations were performed with the snout fully retracted using brass apertures to provide a field size of 18 cm × 18 cm at isocenter. The gantry angle was 180º to allow the water equivalent slabs and 6-well plates to be placed on the patient couch. Thus, the beam pass through the couch, slabs and bottom of the 6 well-plates before reaching the cells. Ionization chamber measurements were performed under the same geometrical conditions using a calibrated parallel plate ionization chamber (PTW 34045). The protective cap was removed from the ionization chamber to minimize the amount of material in front of its sensitive volume. To reproduce the experimental conditions we cut the bottom of three 6 well plates to use in front of the ionization chamber. Irradiations were performed in the same position in respect to the couch and using the same slabs to minimize uncertainties due to material heterogeneity in the couch and slabs. Dose-weighted LET were calculated using a validated Monte Carlo model of the beam line in which the cell lines were exposed.

#### *C-ions*

The C-ion irradiations were performed at HIMAC (Chiba, Japan) with unmodulated beams of energy 290 MeV/nucleon. Water-equivalent energy degraders were used to achieve the dose-weighted LET value of 60.5 keV/µm while the 13.5 keV/µm LET was irradiated in air, with only the small thickness of the acrylic flask or 6-well plate containing the cells modulating the beam. The LET values were calculated as summarized in Kanai et al., 1997.23 All LET values chosen correspond to depths in water that are proximal to the Bragg peak, which were selected so as to minimize the setup uncertainty associated with the high dose and LET gradients in the region of the C-ion Bragg peak. All irradiations were performed at room temperature and all control groups followed the same experimental process minus the radiation exposure.

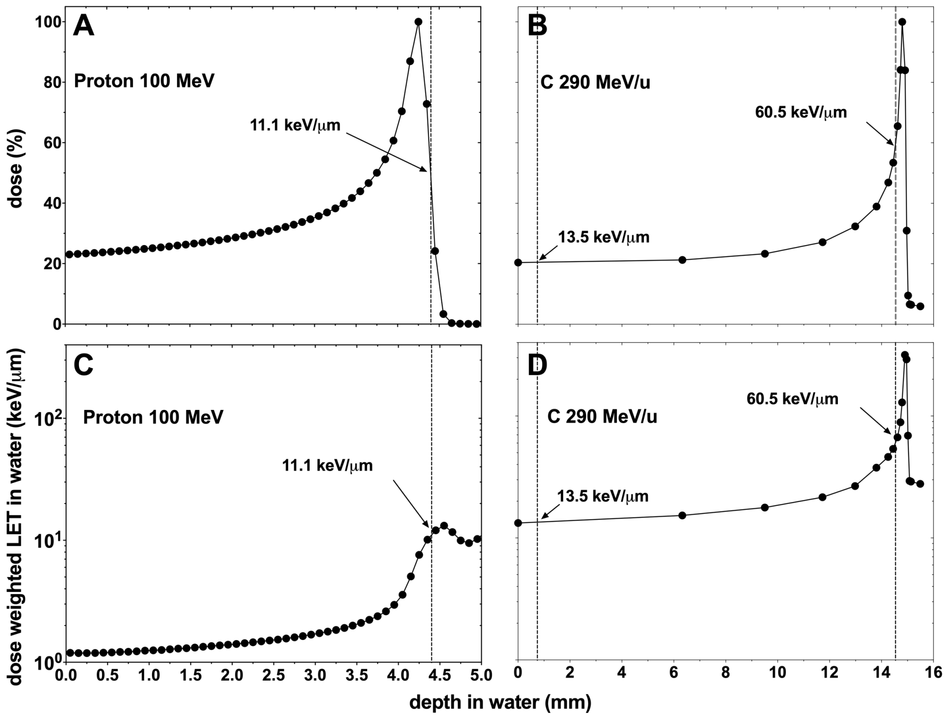


Supplemental Figure 1: Irradiation conditions for the proton and C-ion irradiations. Percentage depth dose for protons (A) and C-ions (B). Dose-weighted LET in water for protons (C) and C-ions (D). Depths at which our cell lines were exposed are shown by dashed lines.

**Supplementary Note 3: Details of clonogenic assay methods**

Cells were seeded into 6-well plates or T12.5 flasks 24 hours before irradiation. After irradiation, the cells were incubated for 8-14 days (see Supplemental Table 3) before being fixed and stained with 0.5% crystal violet in ethanol solution. The dishes containing the stained cell were then scanned using an Epson Expression 10000 XL film scanner, and the images were evaluated using in-house ImageJ macros individually calibrated to score the colonies of each cell line. Briefly, these macros work by blurring convolving the images Gaussian kernel whose radius corresponds to the radius of the smallest colony forming unit (CFU) in order to blur the cells within a colony together. This enables contours to drawn around the colonies, with merged colonies being separated via watershedding. Then colonies whose areas exceed the pixel area threshold, which are individually calibrated to correspond to the area of 50 are more cells, are scored as colonies.
To determine the plating efficiency for each experiment, we fit the data to a function of the form:

$$SQ(D) = PE \cdot e^{-\alpha D - \beta D^2},$$

where SQ is the survival quotient, and represents the survival fraction before accounting for the plating efficient, P, while α and β are free parameters in the fit. We then fit all of the normalized data for a given condition, weighted by the variance, to the linear quadratic model to obtain the final fit:

$$SF = e^{-\alpha D - \beta D^2},$$

We then calculate all radiosensitivity metrics based upon this fit of all the data for each experimental condition.

Supplemental Table 3: Cell lines used in this work, their colongenic incubation period.

| Cell Line | Cancer Histology | Clonogenic Incubation Time |
|---|---|---|
| M059K | Glioblastoma | 9-10 days |
| M059J | Glioblastoma | 9-10 days |
| H1299 | Non-small cell lung cancer | 8 days |
| H460 | Large cell lung cancer | 10 days |
| BxPC-3 | Pancreatic Adenocarcenoma | 14 days |
| AsPC-1 | Pancreatic Adenocarcenoma | 14 days |
| PANC-1 | Pancreatic Adenocarcenoma | 14 days |
| Panc 10.05 | Pancreatic Adenocarcenoma | 14 days |
